# Convergent evolution of epigenome recruited DNA repair across the Tree of Life

**DOI:** 10.1101/2024.10.15.618488

**Authors:** J. Grey Monroe, Chaehee Lee, Daniela Quiroz, Mariele Lensink, Satoyo Oya, Matthew Davis, Evan Long, Kevin A. Bird, Alice Pierce, Kehan Zhao, Daniel Runcie

**Affiliations:** Department of Plant Sciences, University of California Davis, Davis CA 95616

## Abstract

Mutations fuel evolution while also causing diseases like cancer. Epigenome-targeted DNA repair can help organisms protect important genomic regions from mutation. However, the adaptive value, mechanistic diversity, and evolution of epigenome-targeted DNA repair systems across the tree of life remain unresolved. Here, we investigated the evolution of histone reader domains fused to the DNA repair protein MSH6 (MutS Homolog 6) across over 4,000 eukaryotes. We uncovered a paradigmatic example of convergent evolution: MSH6 has independently acquired distinct histone reader domains; PWWP (metazoa) and Tudor (plants), previously shown to target histone modifications in active genes in humans (H3K36me3) and Arabidopsis (H3K4me1). Conservation in MSH6 histone reader domains shows signatures of natural selection, particularly for amino acids that bind specific histone modifications. Species that have gained or retained MSH6 histone readers tend to have larger genome sizes, especially marked by significantly more introns in genic regions. These patterns support previous theoretical predictions about the co-evolution of genome architectures and mutation rate heterogeneity. The evolution of epigenome-targeted DNA repair has implications for genome evolution, health, and the mutational origins of genetic diversity across the tree of life.

**Short Summary:** Fusions between histone reader domains and the mismatch repair protein MSH6 have evolved multiple times across Eukaryotes and show evidence of selection, providing mechanistic and theoretical insight into the forces shaping genomic mutation rate heterogeneity.

## Introduction

DNA damage, the precursor to mutations, occurs between 1,000 to 100,000 times daily in each cell ^1–9^. This poses a continual risk of generating mutations that could negatively impact health and fitness. To address this challenge, organisms rely on proteins that detect and repair DNA damage. Whether organisms can preferentially repair and reduce mutation rates in functionally important genomic regions such as coding regions is both a mechanistic and theoretical question critical to understanding mutation rate evolution ^10–13^.

Research in humans, largely from the field of cancer biology, has revealed specific mechanisms that recruit DNA repair proteins to epigenomic marks found in the coding regions of active genes ^14–34^. These consist of DNA repair proteins fused to histone reader domains, which bind to specific histone posttranslational modification (PTMs). For example, the N-terminus of human DNA mismatch repair protein MSH6 (homolog of bacterial and archaeal MutS^35^) is fused to a PWWP histone reader domain that binds to H3K36me3 (**Figure 1a-c**) ^36,37^. H3K36me3, in humans, is highly enriched in the exons of active genes, while MSH6 plays a key role in several DNA repair pathways. Consequently, targeting MSH6 via its PWWP domain to H3K36me3 reduces mutation rates in coding regions ^18,19,29,38,39^. This is especially true and well-studied in somatic tissues where mutations in coding sequences are the primary source of cancer driver mutations ^29,40^.

**Figure 1.**
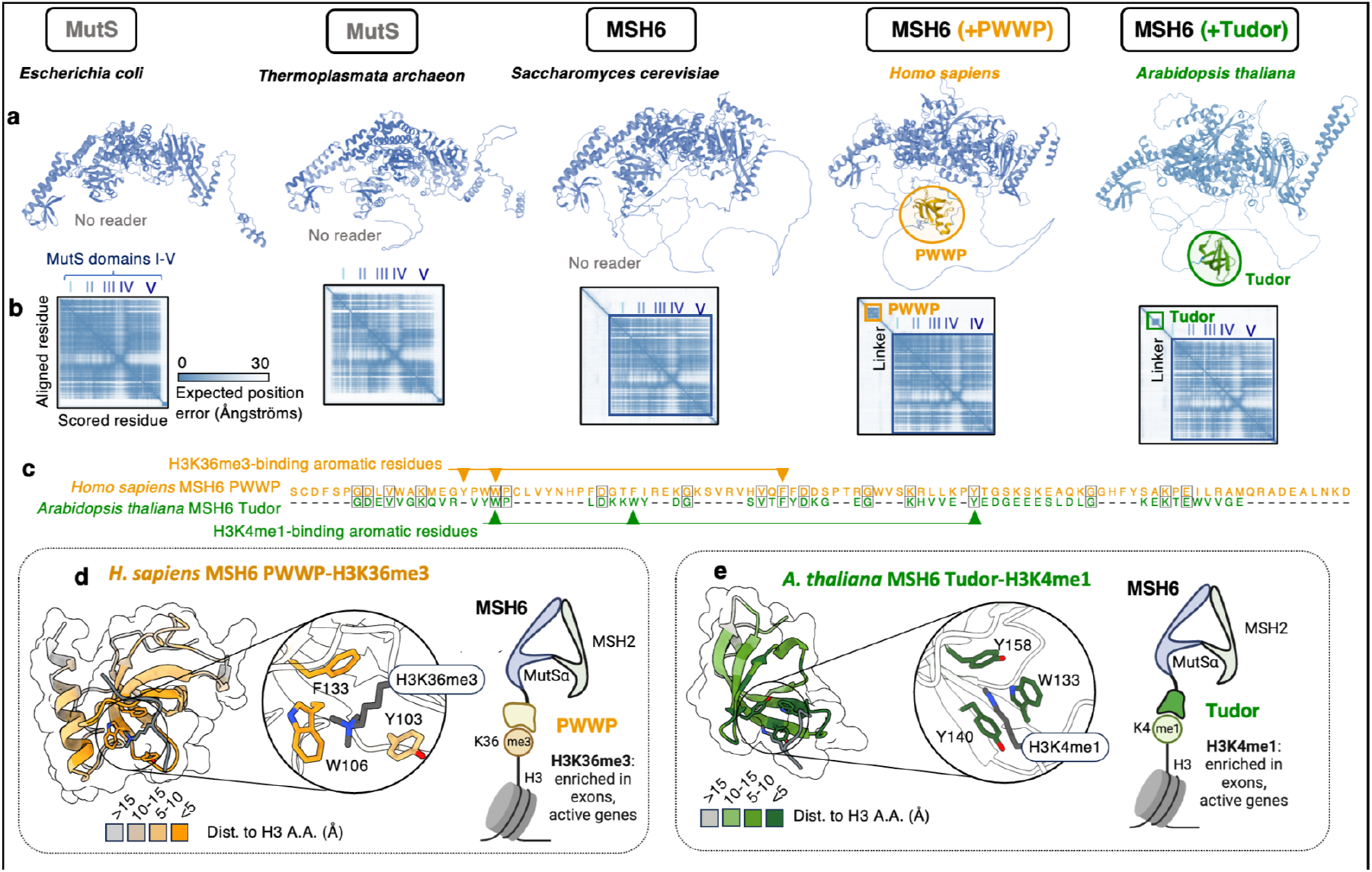
Overview of MSH6 structure and histone reader domains. **a)** Predicted (Alphafold2) structures for MSH6 (MutS homolog), together with **b)** heatmaps showing predicted alignment error, a measure for predicted folding confidence. The “Linker” regions marks the region, predicted to be intrinsically disordered, that separates MutS repair domains from histone reader domains, **c)** Alignment between PWWP and Tudor domains of MSH6, with shared amino acids highlighted by black borders and arrows highlighted experimentally determined amino acids that bind histone PTMs. **d)** Predicted structure of human MSH6 PWWP domain interaction with H3K36me3, highlighting aromatic residues that form binding cage, **e)** Predicted structure of *Arabidopsis* MSH6 Tudor domain interaction with H3K4me1, highlighting aromatic residues that form binding cage.

In plants, H3K4me1 is the most enriched histone PTM, marking coding regions of actively expressed and essential genes ^41–48^. Studies on *de novo* mutation in the model plant *Arabidopsis thaliana* (Arabidopsis hereafter) and rice have found that H3K4me1 exhibits the strongest association with regions of low mutation rates, akin to H3K36me3-associated hypomutation observed in humans ^48–51^. Indeed, mutation rates in Arabidopsis are lower in gene bodies ^49,50,52–60^. Importantly, the N-terminus of MSH6 in Arabidopsis was found to be fused to a Tudor domain that binds H3K4me1, which is distinct from the H3K36me3-binding PWWP domain fused to the human MSH6 (**Figure 1a-d**) ^48,49,58,61^. Experiments have confirmed that genic H3K4me1-associated hypomutation is significantly diminished in the absence of MSH6 in Arabidopsis^48^. This observation aligns with mutation accumulation studies in MSH2 deficient plants (the protein that dimerizes with MSH6 to form the mismatch repair protein complex MutSα ^62,63^), which, like Arabidopsis lacking MSH6, exhibit elevated mutation rates in H3K4me1-rich gene bodies ^56^.

Thus, humans and Arabidopsis have distinct histone readers fused to MSH6, PWWP, and Tudor, respectively, which each bind histone PTMs in genic regions and contribute to lower mutation rates. However, the evolution of these systems across the tree of life remains unknown.

The Tudor and PWWP domains of Arabidopsis and Human MSH6, respectively, are each members of the ‘Royal Family’ of histone readers, which also includes Agenet, Chromo, and MBT domains^64^. These histone readers are found across eukaryotes in numerous proteins (often those involved in chromatin regulatory feedbacks, for example, when fused to histone methyltransferases ^65^). PWWP, Tudor, and other Royal Family histone reader domains are believed to have arisen and diversified from a common eukaryotic ancestor before the emergence of major eukaryotic clades ^64,65^. The differences in binding sites, structure, and amino acid sequence between the MSH6 Tudor and PWWP domains suggest they do not share a common origin in an ancestral MSH6 protein with histone reader functionality (**Figure 1c,d**). Furthermore, the presence of Tudor and PWWP domains in multiple other proteins across Arabidopsis and humans indicates that these domains likely evolved separately before they were incorporated into MSH6 ^64–66^. Thus, the N-terminus fusions of Tudor and PWWP domains to MSH6 may reflect independent evolutionary events, a hypothesis that has yet to be tested with broad phylogenetic analyses.

Here, we investigated histone reader fusions to MSH6 across the eukaryotic tree of life to elucidate their origins, diversity, and potential adaptive value. Specifically, we test for evidence of evolutionary convergence^67^, selection of functional amino acids in the MSH6 histone readers, and co-evolution of MSH6 histone readers with other traits.

## Results & Discussion

### MSH6 independently evolved distinct histone readers in eukaryotes

Our first aim was to determine the evolutionary history and diversity of histone reader fusions to MSH6. We identified putative MSH6 orthologs in 4075 eukaryotic organisms, in which the presence and absence of histone reader could be confidently called, emphasizing the detection and removal of false negatives (**Materials and Methods, Supplemental Table 1**). We observed no cases where MSH6 orthologs showed evidence of containing both PWWP and Tudor domains. The vast majority (99% and 97% of PWWP and Tudor, respectively) of species in which MSH6 histone reader domains were identified showed evidence of histone readers from both *blastp* and *ab initio* domain prediction (**Supplemental Table 2, Supplemental Table 3**).

These data provided the basis to examine the distribution of MSH6 histone reader domains across the eukaryotic tree of life and characterize their evolutionary origins and history (**Figure 2, Figure S1, Figure S2**). We found that fungi, along with most taxa diverging from the last eukaryotic common ancestor, lack PWWP or Tudor domains in their MSH, as also seen in the MutS homologs of bacteria and archaea (**Figure 1a, Figure S3**). This finding is consistent with a reader-less MSH6 being the ancestral state in eukaryotes, supporting independent evolutionary histories of PWWP and Tudor domain fusions to MSH6.

**Figure 2.**
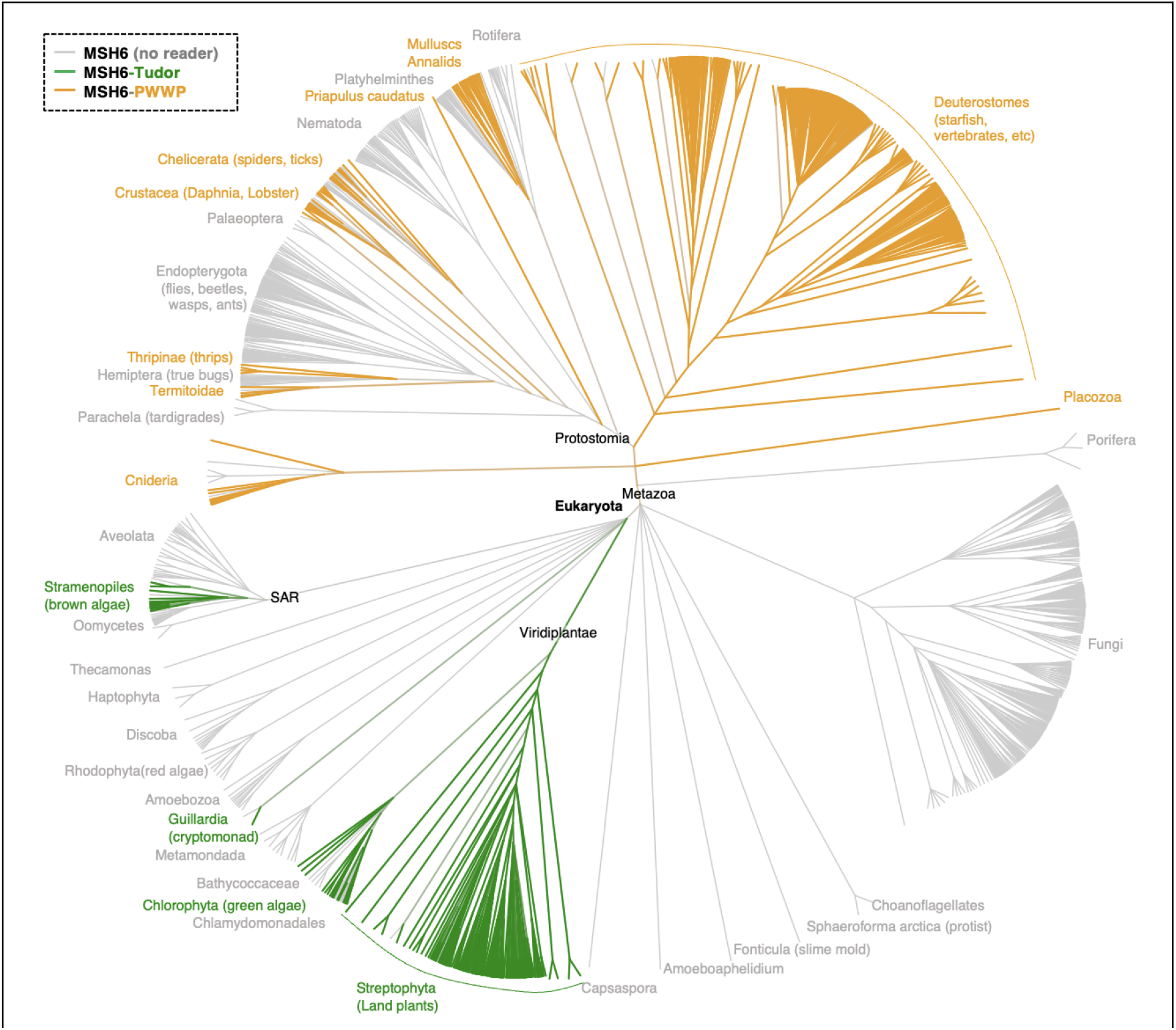
Taxonomic diversity of MSH6 histone reader presence and absence. Tree is based on NCBI taxonomy, showing species for which unambiguous MSH6 orthologs and domain presence/absence could be called. Tree generated with PhyloT, with branches representing taxonomic hierarchy, not time. See Supplemental Table 1 for complete list of species and MSH6 domain calls. For this visualization, fungal genera are represented by a single species. The color of branches indicates proportion of species containing histone reader domains in MSH6 (green = Tudor, orange = PWWP, gray = No reader).

### Metazoa evolved MSH6-PWWP

MSH6-PWWP domain fusions were only found in Eumetazoa, with the distribution of presence/absence suggesting either multiple gains or loss events. While the presence of the PWWP domain is highly conserved in Deuterostomes (98%), major Protostome clades, such as Endopterygota (flies, beetles, wasps, ants) and Nematodes, lack a histone reader (**Figure 2, Figure S1, Figure S2**). Our filtering criteria excluded species with evidence of potential false negatives (MSH6 incorrectly identified as lacking a reader) due to possible genome annotation errors (**Materials and Methods**). Still, individual species that lack evidence of a histone reader in MSH6 are interpreted cautiously until they can be independently verified to be free of assembly or annotation artifacts. Nonetheless, clear phylogenetic patterns exemplified by whole clades lacking any MSH6 reader are likely reliable, along with confirmed reader-less MSH6 orthologs observed in model organisms such as *Drosophila melanogaster, Caenorhabditis elegans*, and *Saccharomyces cerevisiae* (**Figure 1a-b, Figure S1**). These findings are also comparable to broader evolutionary trends in Protosomes, where losses of a number of DNA repair proteins have been observed in various taxa^68,69^.

Determining the ancestral state of MSH6 reader presence in Eumetazoa and whether the variability observed across taxa represents multiple gains versus multiple losses is challenging due to the uncertainty about the relative rate at which these transitions might occur. Losses may be more mutationally likely (i.e., deletion events) than gains (e.g., translocations from another PWWP-containing protein). Ancestral state reconstruction assuming equal rates (ER) suggests multiple gain events, whereas all-rates-different (ARD) models indicate multiple losses (**Figure S1, Figure S2**). The discovery of variation in MSH6 histone reader domains enforces the importance of exercising caution in generalizing experimental findings on mutation rate heterogeneity and DNA repair mechanisms across taxa ^70^. This diversity also has implications for understanding variation in mutation rates and spectra across the Tree of Life ^13^. Once lost, an MSH6 reader may be improbable to be re-gained.

### Plants and their relatives evolved MSH6-Tudor

MSH6 fusions to Tudor domains are unique to land plants and other photosynthetic organisms, including green algae, brown algae, and cryptomonad algae. In land plants (Streptophyta), MSH6-Tudor fusions are highly conserved (96%) (**Figure 2, Figure S1, Figure S2)**. Some green algae (Chlorophyta), along with some diatoms, brown algae (Stramenopiles), and other SAR (Stramenopiles + Alveolata + Rhizaria) lineages lack the Tudor domain (**Figure 2**). It is interesting to note that the species lacking MSH6-Tudor includes *Chlamydomonas reinhardtii*, a model for algal genomics, which also lacks the H3K4me1 enrichment of gene bodies that is otherwise conserved in higher land plants ^44,71^. We also do not observe MSH6-Tudor in any red algae (Rhodophyta). Like in animals, the ancestral state of the MSH6 domain presence is somewhat uncertain, especially given that several groups containing Tudor domains, including Stramenopiles, have potentially gained genes from green algae via ancient endosymbioses ^72,73^. However, the phylogenetic relationship of MSH6 orthologs does not support horizontal gene transfer from green algae: a maximum likelihood tree of MutS domains I-V shows the MSH6 ortholog of Stramenopiles sister to other SAR taxa that lack a Tudor domain (**Figure S4**).

Plants (Viridiplantae) contain a unique MutS ortholog, MSH7, which arose via an ancient duplication of MSH6 ^74–76^. We identified putative MSH7 orthologs in plants and queried whether Tudor domains were observed (**Materials and Methods**). We found no plant MSH7 orthologs containing Tudor domains, suggesting that the Tudor was acquired by MSH6 after the duplication that gave rise to MSH7 (**Figure S6**). or that MSH7 lost the Tudor domain during or after duplication in the common ancestor of plants. Thus, the MSH6-Tudor fusion is probably ancient and predates the separation of the major clades of Viridiplantae, possibly in the common ancestor of Viridiplantae, crytomonads, and SAR.

### Testing for additional fusions of MutS homologs to histone readers

We used *ab initio* domain prediction to investigate whether additional, previously undescribed, histone reader fusions have occurred with MSH6. This analysis yielded no known candidate histone reader domains beyond Tudor and PWWP (**Supplementary Table 3**). We further expanded our search to other potential MutS homologs, and while we did identify a number of intriguing fusions to non-MutS domains, we found none with obvious histone readers.

For example, we identified a fusion between MSH3 and N-acetylglucosaminyl transferase component (Gpi1) (e.g., UniProt: G3Y5U7) shared across *Aspergillus* fungi, with unknown functional importance. We also found a fusion of MSH6 with ESCO acetyltransferase and Zinc-Finger binding domains in the Ustilaginomycetes (smut fungi) (**Figure S5**). These domains are not known to be histone readers *per se*; however, ESCO proteins have been previously described to bind chromatin by DNA-binding of a charged N-terminus intrinsically disordered region ^77^. While not histone readers, these findings are a reminder of the potential for undiscovered fusions between chromatin interacting domains and other proteins involved in DNA repair, a potentially valuable direction for future research.

In summary, we find compelling evidence of convergent evolution in the origins of distinct histone-binding domains, specifically to MSH6 in eukaryotes^67^. These fusions of histone readers to MSH6 provide a unique opportunity to study the evolutionary dynamics of epigenome-recruited DNA repair systems at a Tree of Life scale.

### Conservation of domain architectures and sequence

We next set out to further test if the PWWP and Tudor domain fusions to MSH6 exhibit evidence of being subject to natural selection. Patterns of conservation can also provide mechanistic insight into the adaptive value and biochemical function of MSH6-associated histone readers.

#### Constraint on linker region

In both human and Arabidopsis, the MutS repair domains are separated from the histone reader by a (MobiDB-lite annotated) intrinsically disordered “linker” region (**Figure 1a-b, Figure 3a**). To better understand this region, we examined its sequence composition and length across all species containing the respective domains (**Figure 3a**). Regardless of histone reader (PWWP or Tudor), we found that MSH6 linker regions were significantly (p<2×10^-16^) enriched for the charged amino acids lysine (K), aspartate (D), and glutamate (E), consistent with a role in promoting disordered protein regions and potentially other molecular interactions^78^. As expected, these linker regions were >36 times more likely to be predicted as intrinsically disordered than histone reader or MutS-containing protein regions *in silico*^*79*^ (Z=782.5, p<2×10^-16^).

**Figure 3.**
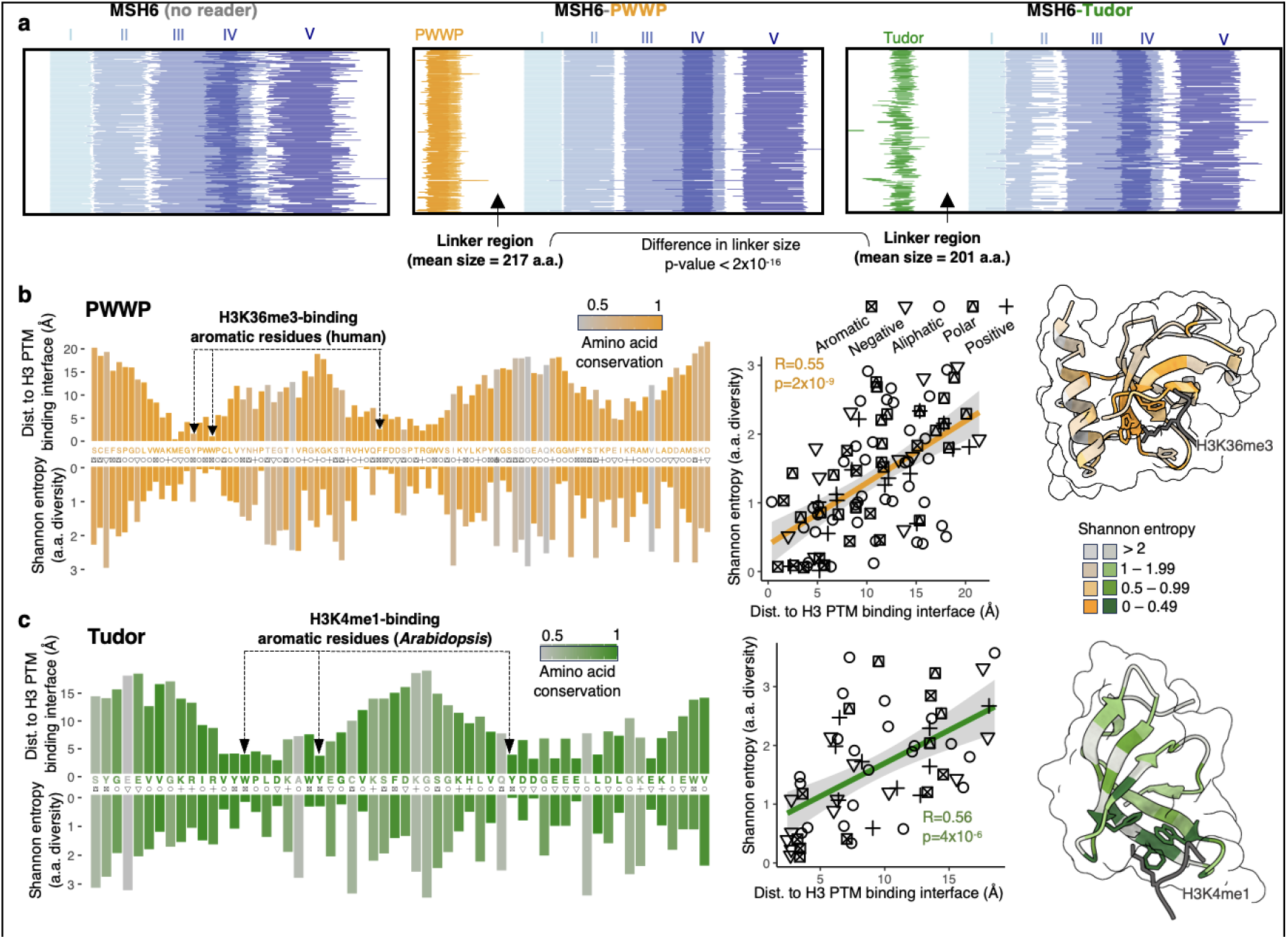
Conservation in MSH6 domain architectures and histone readers. **a)** MSH6 domain architectures (InterProScan) for MSH6 protein (200 random species with and without histone reader domains, aligned to the MutS1 domain of MSH6. Relationship between distance to H3 amino acids and amino acid Shannon entropy across all species, colored by conservation (proportion AA = consensus) for **b)** PWWP domains and **c)** Tudor domains. Consensus amino acids with corresponding functional class shown. Amino acid conservation is equal to the frequency of the most frequently observed amino acid. Distances: Tudor + H3K4me1 approximated by superimposition of domains to reference PDB 7*DE*9; PWWP + H3K36me3 with reference PDB 5*iu*.

We also observed a notable degree of conservation in linker sequence lengths (**Figure 3a**). It is difficult to establish a strong null model with which to compare disordered region length variation against neutrality. However, we did note that Tudor domains are significantly closer (shorter linker region) to MutS domain I than PWWP (Tudor mean = 201 a.a., PWWP mean = 217 a.a, t = 13.527, p-value < 2.2×10^-16^, **Figure 3a**). This is interesting given that the Arabidopsis MSH6 Tudor and human PWWP domains bind to methylated lysines on the 4th (most distal to histone/DNA) and 36th (more proximate) lysines, respectively. The length of the disordered and potentially flexible linker region may be constrained to tether MSH6 at an optimal distance to chromatin, a hypothesis necessitating further experimental testing.

#### Constraint on histone interacting domain amino acids

We then investigated patterns of evolutionary constraint on the reader domains themselves. If the histone PTM binding affinities are subject to selection, we would expect to observe the conservation of amino acids involved in interactions with the H3 peptides, as seen in the evolution of sites involved in protein-protein interactions^80^. We compared MSH6 Tudor and PWWP domains across all species. For each amino acid position within the domains, we identified the consensus amino acid (the most common residue) and calculated both the conservation (the frequency of the consensus amino acid) and Shannon entropy (a measure of amino acid diversity at each position).

For both Tudor and PWWP domains, amino acids involved in H3 PTM binding were among the most conserved residues, consistent with purifying selection on histone reader binding affinities **(Figure 3b-c**). We estimated the distance between amino acids and the H3 peptide by superimposing predicted structures to experimentally determined PWWP and Tudor structures that captured H3 PTM binding, H3K36me3 and H3K4me1, respectively (**Materials and Methods**). We found that amino acids predicted to be in close proximity to the H3 peptides are significantly more conserved, both in PWWP and Tudor reader domains (PWWP: R=0.55, p=2×10^-9^; Tudor: R=0.56, p=4×10^-6^). **(Figure 3b-c**). Thus, evidence of purifying selection was enriched at the histone-binding interface. Several distal amino acids were also highly conserved. This conservation may reflect roles in proper folding or other interactions, such as between PWWP and DNA directly ^81^.

In summary, evolutionary conservation observed in MSH6 domain architectures, as well as Tudor and PWWP domains themselves, exhibit signatures of selection, especially for histone binding.

### Species with MSH6 histone readers show distinct genomic traits

The preceding analyses revealed convergent evolution and conservation of MSH6 histone readers (**Figure 3**). However, not all Eukaryotes gained (or retained) MSH6 histone readers (**Figure 2, Figure S3, Figure S4**). To study this variability further, we tested for correlated evolution of MSH6 reader presence/absence with genomic architectures and other traits toward understanding what might be behind the observed phylogenetic patterns.

#### Contrasting genomic traits

Theory and experimental work have observed that hypermutators and changes in mutation biases can be favored by selection under certain conditions ^82–84^. Whether this can help explain variability in MSH6 reader presence in eukaryotes is a compelling, albeit challenging, question to answer. Previous theory based on the drift-barrier hypothesis predicts that the adaptive value of promoting DNA repair in coding regions may be contingent on genome architectures^13^. To test this, we compared a suite of genomic traits between species (**Figure 4, Figure S7, Table S4**). We found that species with MSH6 fused to histone readers have (more than an order of magnitude) larger genomes. Previous investigations have described substantial evolutionary variation in intron/exon architectures across eukaryotes ^85,86^. After accounting for phylogeny, we found significant co-evolution of MSH6 readers with genome size, marked especially by greater intron content, both in size and number per gene (**Figure 4b-c, Figure S7a**). This was also the case when only Protosome animals were examined, where the loss of MSH6-PWWP may have occurred (**Figure S7b)**. These observations are consistent with the hypothesis proposed by Lynch et al. (2023)^13^ that “the benefits of targeting DNA repair to coding regions may be greatest for species with a large fraction of non-coding genome content.”

**Figure 4.**
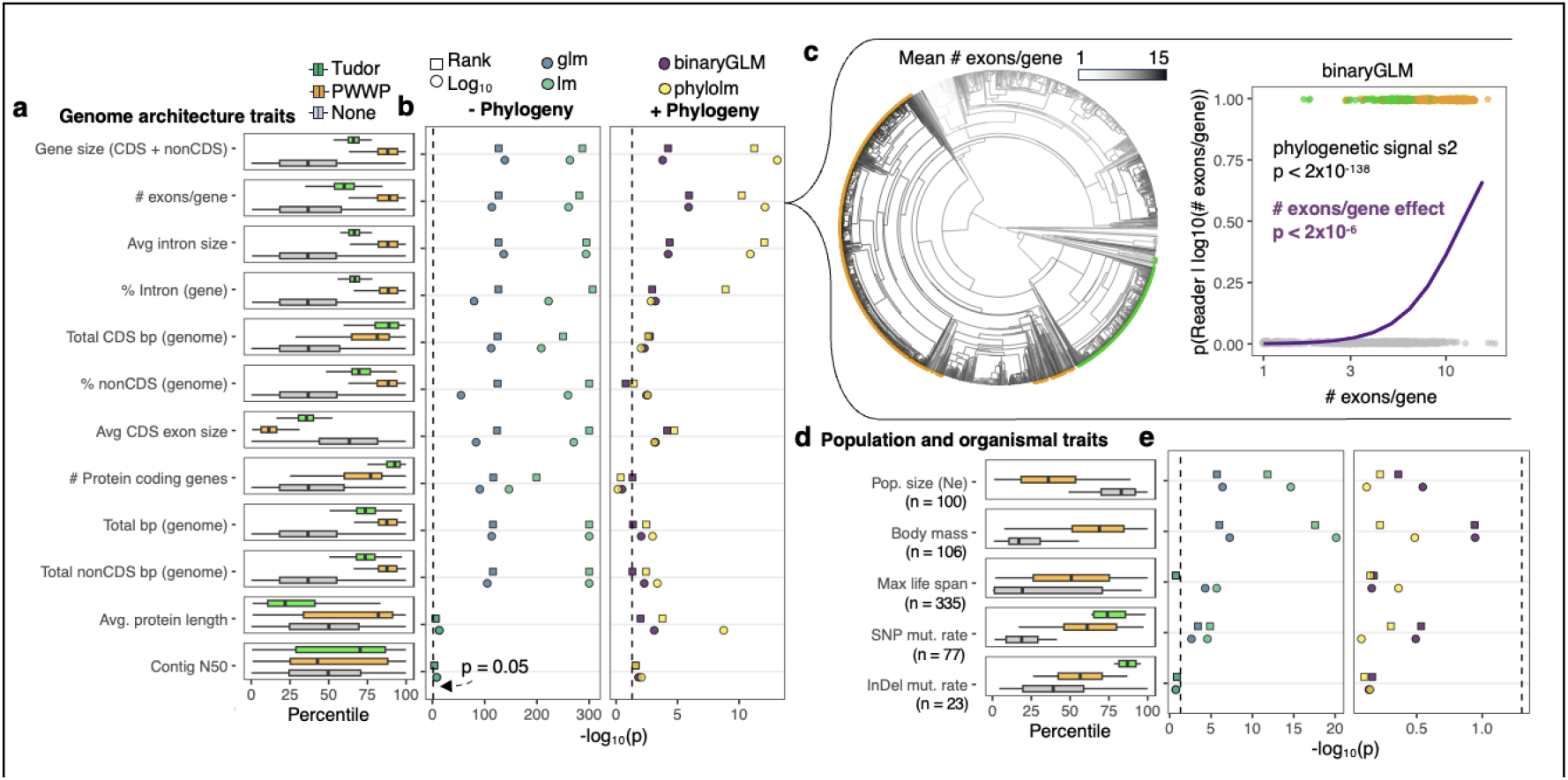
Species with histone readers fused to MSH6 differ in genomic and other traits. **a)** Summary of genomic traits among species with MSH6-Tudor, -PWWP, or no reader, scaled to percentile. Boxplots: box = median± interquartile range (IQR), whiskers = 1.5 x IQR. **b)** Summary of contrasts: binomial generalized linear model and linear model (left), phylogenetic binary regression with phylolm::phy1olm and with ape::binaryPGLMM (right) using TimeTree V5 phylogeny (Kumar et al. 2022). Analyses for log_10_ transformed and rank values were tested. Binomial tests compare species with either Tudor or PWWP vs species with no reader, **c)** Example of phylogenetic trend in in mean #exons/gene in relation to the presence of MSH6 reader. Left: ancestral reconstruction of mean logw(#exons/gene) across eukaryotes, with tips colored by presence of Tudor (green) and PWWP (orange) domains in MSH6. Right: results from binomial phylogenetic regression, **d)** Population and other traits across species as described in **(a)**. Number of species with these data are indicated by (n = X). Estimated body size and effective population size from Buffalo 2021. Lifespan from Tacutu et al. 2018. Mutation data from Wang & Obbard 2023. e) Statistical contrasts as described in **(b)**.

For species where data were available^87,88^, we also compared several metrics relevant to life history and population genetics. Species without MSH6 histone reader domains have smaller body sizes, greater estimated population sizes, and experience significantly lower germline SNP mutation rates (**Figure 4d**). While interesting, these associations were not significant after accounting for phylogeny. Measurements of these characteristics across more species with a broader taxonomic range may be needed for more robust tests for co-evolution with reader domains in phylogenetic contexts.

In summary, we observed significant co-evolution with genome features like genome size and intron number per gene and found differences in population and organismal characteristics in species with and without reader fusions to MSH6. Whether these reflect a cause or consequence of epigenome-recruited repair or instead reflect correlated responses to shared evolutionary forces remains an open question. Still, these results highlight interesting connections between DNA repair systems and the evolution of genome architectures. These patterns are consistent with the selective benefits and costs of such systems being contingent on factors like non-coding genome size, as previously hypothesized^13^. More sampling of traits and mutation rates, coupled with theoretical analyses, are needed to integrate these findings through the models of the drift-barrier hypothesis to better determine what evolutionary processes underlie these associations^10–13,49,89^.

These findings inspire a number of outstanding questions: How conserved are epigenomes across the tree of life? Are there yet-to-be-discovered mechanisms of epigenome-recruited DNA repair systems? Are there biological or evolutionary costs associated with epigenome-recruited mismatch repair? What is the relative importance of these mechanisms for somatic versus germline mutation in different organisms? Do the distributions of fitness effects between coding and non-coding sequences differ among organisms? Finally, what is the extent of mutation rate heterogeneity across different organisms’s genomes? These and other questions, in light of the findings presented here, provide a roadmap for future empirical and theoretical work.

## Conclusions

The recruitment of DNA mismatch repair by H3K36me3 (MSH6-PWWP) in humans and by H3K4me1 in Arabidopsis (MSH6-Tudor), now investigated at a tree of life scale, reveals a compelling example of evolutionary convergence^67^. Previous experiments support the role of these mechanisms in recruiting mismatch repair to functionally important genomic regions, at least in Humans and Arabidopsis, with consequences on mutation rates ^17,37,38,48,56^. The phylogenetic distribution of PWWP and Tudor fusions to MSH6 support their independent evolutionary origins, a hallmark of evolution by natural selection. Patterns of evolutionary conservation reveal selection for histone binding and potentially on the regions that link reader domains with MutS repair domains, providing hypotheses about biochemical functions. We also find evidence of clades that never gained or have lost MSH6 histone readers, especially of PWWP in major clades with Protosome animals, coupled with differences in genome architectures and other characteristics. This co-evolution may provide insight into the evolutionary forces acting on these mechanisms, reflecting their costs and benefits under different conditions. In summary, the convergent evolution of epigenome targetting in MSH6 suggests conditionally adaptive mechanisms, inspiring further questions about the mutational consequences and evolution of DNA repair systems across the tree of life.

## Materials and Methods

### Defining MSH6 orthologs

We downloaded all annotated (both NCBI RefSeq and GenBank) public eukaryotic genomes and proteomes on NBCI (4,938 species) on August 9, 2023 (**Figure S8**). For each proteome, we queried using the protein sequence of human MSH6 (*blastp* with eval threshold of 0.01). Each of the resulting hits was blasted back against the human proteome. Proteins were considered potential orthologs of human MSH6 if the top hit in the human proteome was human MSH6. We thus removed proteins that show evidence of belonging to other members of the MSH protein family, though these were retained for downstream analysis to test for the presence of histone readers. We also performed the same reciprocal blastp analysis with Arabidopsis MSH6. Proteins were considered MSH6 only if the top hit of the reverse blast was the Arabidopsis MSH6 protein. These predicted MSH6 orthologs for each species were then defined as those that were classified as orthologs of both human and Arabidopsis MSH6. Finally, to further confirm MSH6 orthology, we ran *ab initio* of each predicted MSH6 ortholog using InterProScan with all available databases^90^. The final set of MSH6 orthologs was determined by requiring them to be called MSH6-specific orthologs by both reciprocal *blastp* queries and also contain *ab initio* annotated MutS domains. This removed potential false positives from downstream analysis in which non-MSH proteins containing PWWP or Tudor domains would be incorrectly called. Because genome annotation errors can lead to false negatives of MSH6 presence in a species, we ignored all species lacking a definitive MSH6 ortholog in all downstream analyses.

### Identifying Tudor and PWWP domains in MSH6 orthologs

For each species proteome, we queried for the PWWP domain of the human MSH6 protein (**Figure S8**). We also queried for the Tudor domain of the Arabidopsis MSH6 protein. From these results, we annotated MSH6 orthologs as containing Tudor and PWWP domains if the same protein was detected by MSH6 blastp searches (preceding step) and significant (e score) matches for specific queries of PWWP and Tudor domains. To further improve the annotation of Tudor and PWWP domains, we analyzed the results of *ab initio* domain prediction from InterProScan with “CDD,” “Pfam,” “SMART,” and “ProSiteProfiles” databases. MSH6 orthologs were thus annotated as containing a Tudor domain if they contained a significant match to the Tudor domain query or if they were annotated as having a Tudor domain by InterProScan. Likewise, MSH6 orthologs were thus annotated as containing a PWWP domain if they contained a significant match to the PWWP domain query or if they were annotated as having a PWWP domain by InterProScan. Species were classified as containing a Tudor domain or a PWWP domain in MSH6 if any of their MSH6 orthologs contained such domains. This approach was conservative in that it prioritized against calling false negatives. That is, to preclude misclassifying species as lacking a histone reader when, in reality, one is present. For example, in a polyploid species, we might find multiple putative MSH6 orthologs, which vary in the presence/absence of a reader domain. If at least one MSH6 ortholog contained a reader domain, it was classified as having that domain at the species level.

### Removing species with evidence of genome annotation errors

We considered several possible scenarios from which genome annotation errors could cause false negatives in identifying reader domains in MSH6 orthologs (**Figure S8**).

First, genome annotations could lead to incorrect splits of MSH6 into several genes. In this case, the MSH6 gene may be found adjacent to a “gene” that contains a Tudor domain or PWWP domain. To identify such cases, we compared the location of MSH6 orthologs to results from querying (*blastp*) Tudor and PWWP domains. Cases in which a neighboring gene (within 5kb) contained a Tudor or PWWP domain were classified as ambiguous and removed from subsequent analyses.

Second, genes could be incompletely annotated such that transcribed regions (e.g., whole exons) are excluded from the predicted gene model. We thus used *tblastn* to query Tudor and PWWP domains on the entire genome, which were translated into all reading frames of each genome. Species in which a sequence matching a Tudor or PWWP domain within 5 kb of an MSH6 ortholog were classified as ambiguous and removed from subsequent analyses.

Finally, we considered the possibility that MSH6 orthologs with histone reader domains may be missed from annotation entirely. To identify and remove such cases, we used *tblastn* to query MSH6 protein sequences from human and Arabidopsis orthologs. Cases where potential unannotated orthologs (matching MSH6 > 200aa, to exclude matches to potential reader domain itself, such as in other proteins which contain these domains) were defined as cases where a *tblastn* query for Tudor or PWWP domain was also found within 5 kb of MSH6 tblastn result. For such cases, the species’ domain call was classified as ambiguous and excluded from downstream analyses.

Despite these steps, we could not preclude other sources of misannotation that might lead to false negatives for domain presence in a single species. For example, fragmentation during genome assembly could lead to edge cases where the repair domain (i.e., MutS domains) of MSH6 and its reader domain (if present) were assembled on different scaffolds or contigs, making it impossible to accurately control against the subsequent effects of misannotation. Similar issues might arise from miss-assemblies that cause false separation from MSH6 repair domains from reader domains. Thus, we interpret single species lacking a reader domain, especially when nested within a clade in which closely related species that contain such a domain, with caution until they can be validated.

### MSH6 homologs in non-eukaryotes

We first examined the structure and domain architectures of MutS homologs in Escherichia coli (strain K12) (Uniprot: P23909) and *Thermoplasmatales archaeon* (Uniprot: A0A256YTS1), which showed no reader domains (in structure nor in Interproscan annotations). We queried, with *blastp*, the entire NCBI Archaeal protein database for the Arabidopsis and Human MSH6 proteins and checked for any that aligned to Tudor or PWWP regions of MSH6. We also queried Archaeal proteomes for the Tudor and PWWP domains directly, which yielded no hits. We repeated this specifically for Asgard Archaea, described as the closest relatives to Eukaryotes ^91^.

### Ancestral state reconstruction

To estimate the likely ancestral states, we used the ace function from *ape* in R, which estimates marginal ancestral states for a discrete trait (in this case, MSH6 reader domain state) with the re-rooting method described by Yang et al. (1995)^92^. For this, we used the time-calibrated eukaryotic species tree (2004 species) available from TimeTree v5 ^93^, setting the edge.length + 1×10^-5^ to account for 116 branches with length=0. We compared ancestral states estimated with “equal rates” (ER) and “all-rates-different” (ARD) models. We also estimated ancestral states for “PWWP” vs. “Tudor” vs. “None,” as well as “PWWP” vs. “None” and “Tudor” vs. “None” separately.

### Natural variation in Tudor and PWWP domains

Tudor and PWWP domains (± 5aa from respective *blastp* match) of MSH6 from all species were extracted from protein fasta files using blastp hits to define boundaries of domains. The resulting unique sequences were aligned with MUSCLE with the msa function from the R package *msa*. We used the Arabidopsis Tudor and human PWWP as reference frames to define alignment positions along the domains and thus ignored insertions relative to these. From these alignments, we calculated the frequency of each amino acid at each position and the corresponding Shannon entropy *H(X)* as

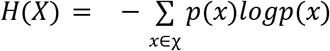

Where *p(x)* represents the frequency of an amino acid *x* from alphabet *χ* (20 amino acids and gap) at position *X*. We also calculated conservation at each position as the proportion of the most frequent amino acid, with these amino acids representing, together, the consensus sequence across the domain.

### Modeling Structure of Tudor and PWWP domains interface with histone PTMs

Intrinsically disordered regions (IDRs) in all MSH6 proteins were predicted with ALBATROSS ^79^. To examine the size, sequence composition, and IDR enrichment of the linker sequence separating MutS domains from histone readers, we used results from InterProScan to identify positions of well-annotated protein domains (one each of MutS I through V, Tudor, and PWWP, if present). Amino acid enrichment in linker regions was estimated by comparing amino acid counts within linker regions across species with the expectation of uniformity in amino acid frequencies. IDR enrichment was estimated by comparing the frequency of overlaps between linkers and predicted IDRs with overlaps between repair or reader domains with predicted IDRs. Given the disparities in species numbers with each domain architecture, we randomly selected 200 species with each domain architecture to visualize (Figure 3a). These were plotted with domain positions centered around the start of MutS domain I. We compared the estimated size of the linker sequence between all PWWP and Tudor domain-containing species by measuring the length between the InterProScan annotated reader and MutSI domains.

Next, we approximated the distance between reader domain amino acids and H3 peptides. To obtain these estimates, we superimposed the Human MSH6 PWWP domain predicted by Alphafold ^94^ onto PDB *5ciu* ^*95*^, the experimental structure of H3K36me3 bound by the DNMT3B PWWP domain, using the Pairwise Structure Alignment tool from RCSB with the TM-align method (https://www.rcsb.org/alignment). We superimposed the Arabidopsis MSH6 Tudor domain to PDB *7DE9*^*45*^, the experimental structure of H3K4me1 bound by the PDS5C(RDM15) Tudor domain. From the resulting structural alignments, we estimated the distance of each amino acid in the histone reader domain to the corresponding H3 peptide chain by calculating the minimum Euclidean distance between atoms in each amino acid to atoms in the H3 peptide. We compared the relationship between the estimated distances of each amino acid with their evolutionary constraint (Shannon entropy). Similar results were found when we compared distance with conservation.

### Comparisons between species with and without reader domains in MSH6

For each species’ genome and its corresponding annotations (averaging summary statistics across annotations when multiple were available), we estimated the total genome size by the assembly length in base pairs and calculated the total base pairs of protein-coding sequence, total base pairs of non-protein-coding sequences, the average number of coding exons per protein-coding gene, average length of coding exons, average length of introns per gene, number of introns, average proportion of introns in gene bodies, and average length of proteins. To validate genome size, we compared genome assembly size with previously reported C-values ^96^ and found high concordance (r=0.97, p<2×10^-16^). As a further control against assembly artifacts influencing results, we also tested contig N50 values as reported in NCBI in the same manner as other variables. Other traits: Lifespan was compiled by the AnAge database ^97^. Germline mutation rates previously compiled by Wang & Obbard (2023)^88^. Body size (“log10_body_mass_pred”) and population size (“log10_popsize”) estimates were generated by

Buffalo (2021)^87^. Of these, we used both log_10_ transformed and ranked values to account for non-normality. We first performed binomial generalized linear models contrasting species with and without Tudor or PWWP histone reader domains in R with glm(family=“ binomial”). We then performed phylogenetically constrained contrasts using the time-calibrated eukaryotic species tree (2434 species) reported from TimeTree v5 ^93^, setting the edge.length + 1×10^-5^ to account for 116 branches with length=0. We compared histone reader presence/absence classification as a function of the above metrics with the *phyloglm* function (btol = 1000, method=“logistic_MPLE”, log.alpha.bound=0) from the R package *phylolm* V2.6.2 and the binaryPGLMM (default settings) function from the *ape* V5.7.1 package ^98,99^. We also modeled the relationship between readers and traits with comparative.data (vcv=TRUE, vcv.dim=2) and pgls functions (lambda=‘ML’) from the *geiger* V2.0.11 package in R and *phylolm* function from the *phylolm package* (model=“kappa”).

## Acknowledgments

We thank Dan Sloan, Detlef Weigel, and Michael Lynch for valuable discussions of this work. The Monroe Lab (J.G.M) is supported by FFAR grant ICRC20-0000000014, USDA-NIFA grant 108681-Z5327202, a UC Davis STAIR grant, NSF Grants 2317191 and 2338236, and CPRB grant HG-2022-35. The research was conducted at the University of California Davis, which is located on land that was the home of the Patwin people for thousands of years (https://diversity.ucdavis.edu/land-acknowledgement-statement).

## Data availability

Genomes and annotations analyzed here are available on NCBI and listed in Table S1. A summary of data used to classify MSH6 histone readers is found in Table S2. Interproscan results for putative MutS homologs are found in Table S3. Species-level genome and trait information is found in Table S4.

## Code availability

Code for this research is maintained at github.com/greymonroe/MSH6_evolution

## Supplemental Materials

## Supplemental tables

Table S4. Species-level genomic and other characteristics

Table S3. Interproscan results

Table S2. Evidence for the presence/absence of MSH6 histone reader domain

Table S1. Summary of presence/absence of histone reader domains and genomic traits across species

## Supplemental Figures

Figure S1. MSH6 tree with branch lengths

Figure S2. Ancestral state reconstruction

Figure S3. Archaea MSH6

Figure S4. MutS domain tree

Figure S5. MSH3 fusion with domain in fungi

Figure S6. MSH7 examples in plants

Figure S7. Traits in relation to reader presence

Figure S8. Methods for classifying MSH6 reader presence

**Figure S1.**
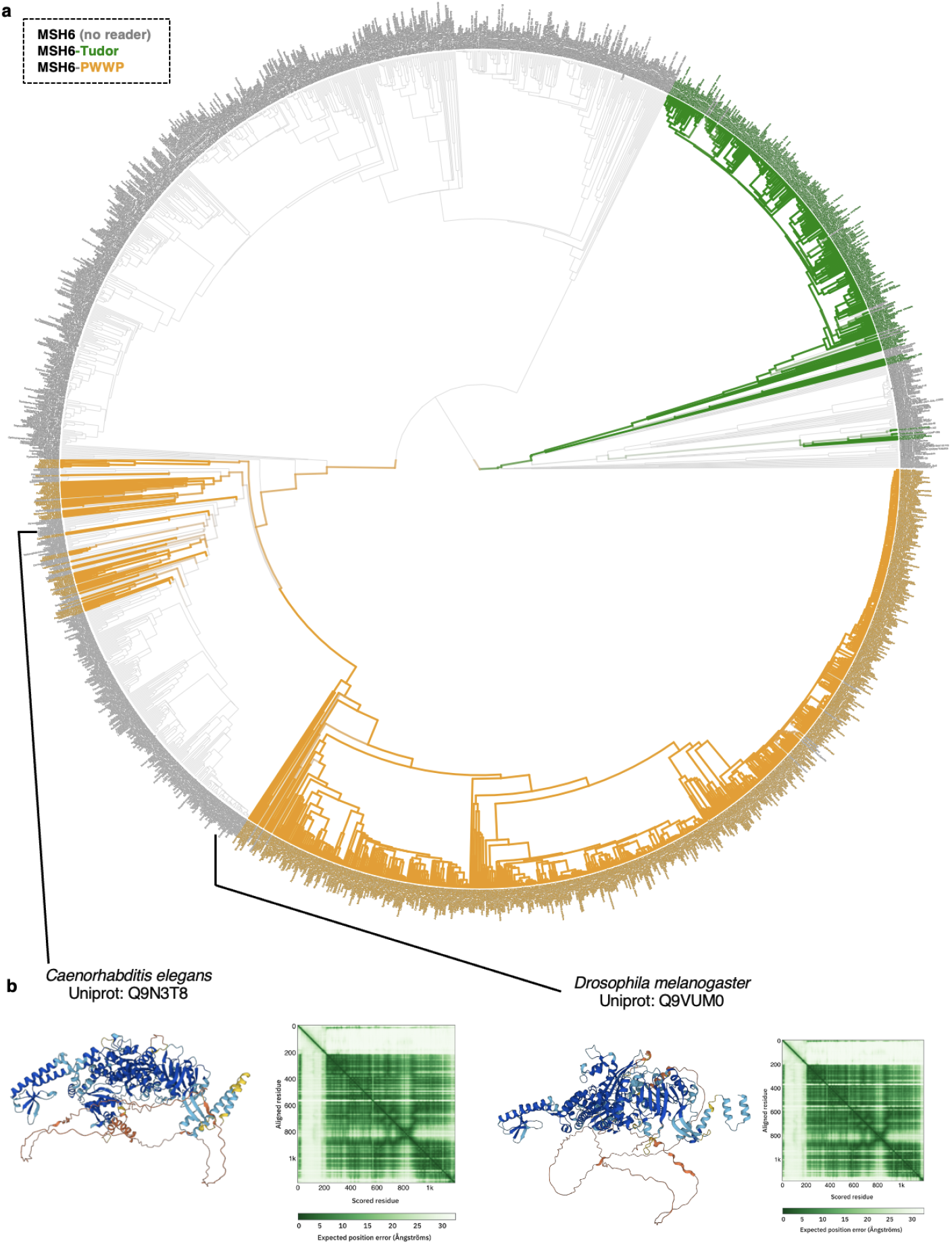
Phylogeny of of Eukaryotes with branch lengths. **a)** Tree from TimeTree v5 with branches and tips colored by presence of MSH6 reader domains. Contains 2004 species. See Supplemental Table 1 for complete list of species, **b)** Illustrative examples of MSH6 lacking histone reader domain in Metazoa. Alphafold predicted structures of MSH6 in the model organisms, *Drosophila melanogaster* and *Caenorhabditis elegans*.

**Figure S2.**
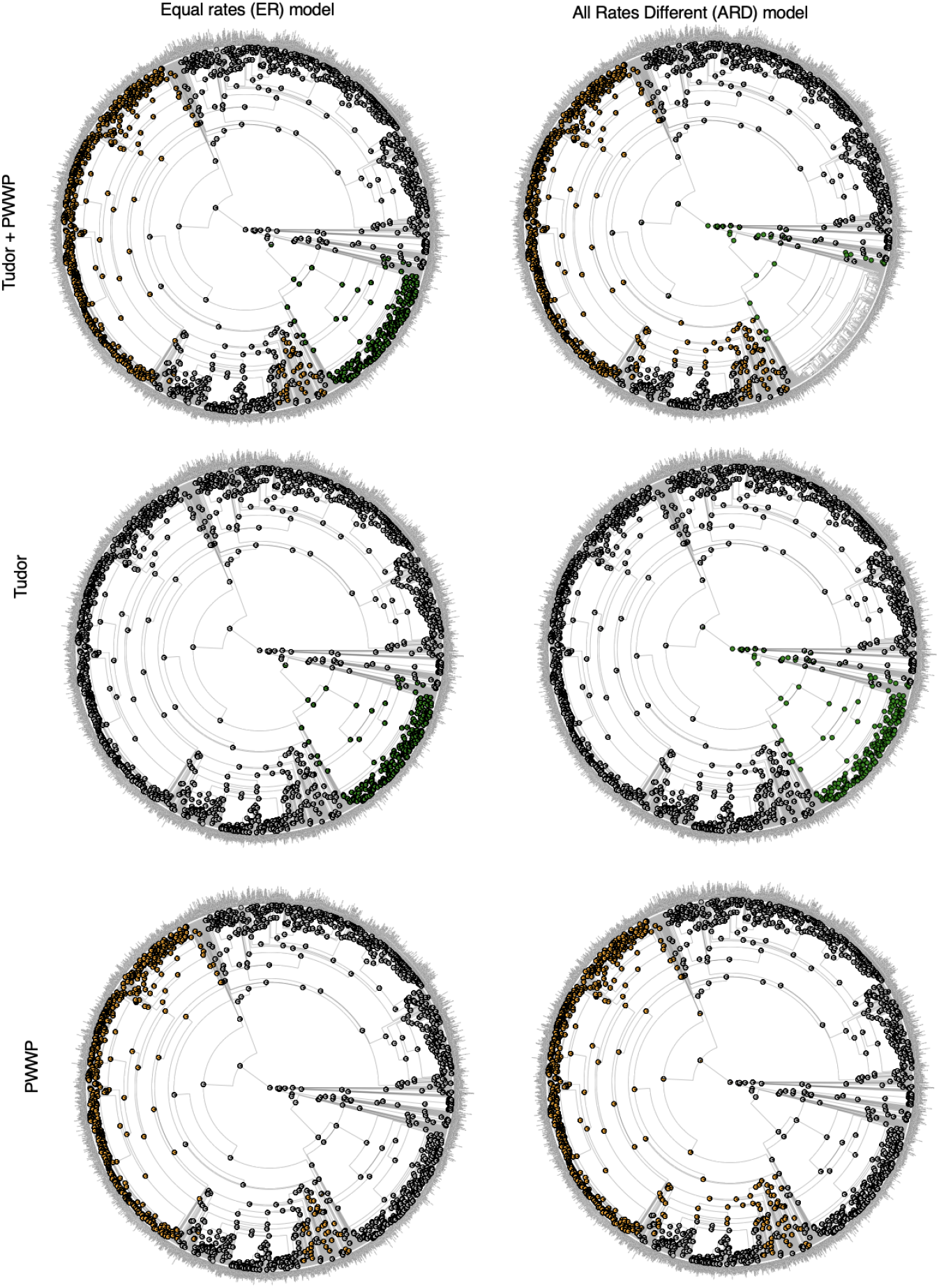
Ancestral reconstruction of MSH6 reader domain presence and absence. Generated with ace function from ape in R. Left shows results from ace(reader ,tree,type=“discrete”,model=“ER”), right shows ace(reader ,tree,type=“discrete”,model=“ARD”)· Top panels show reconstruction of all readers, middle shows Tudor alone, and bottom shows PWWP. Pie plots at internal nodes show the posterior probability of ancestral state.

**Figure S3.**
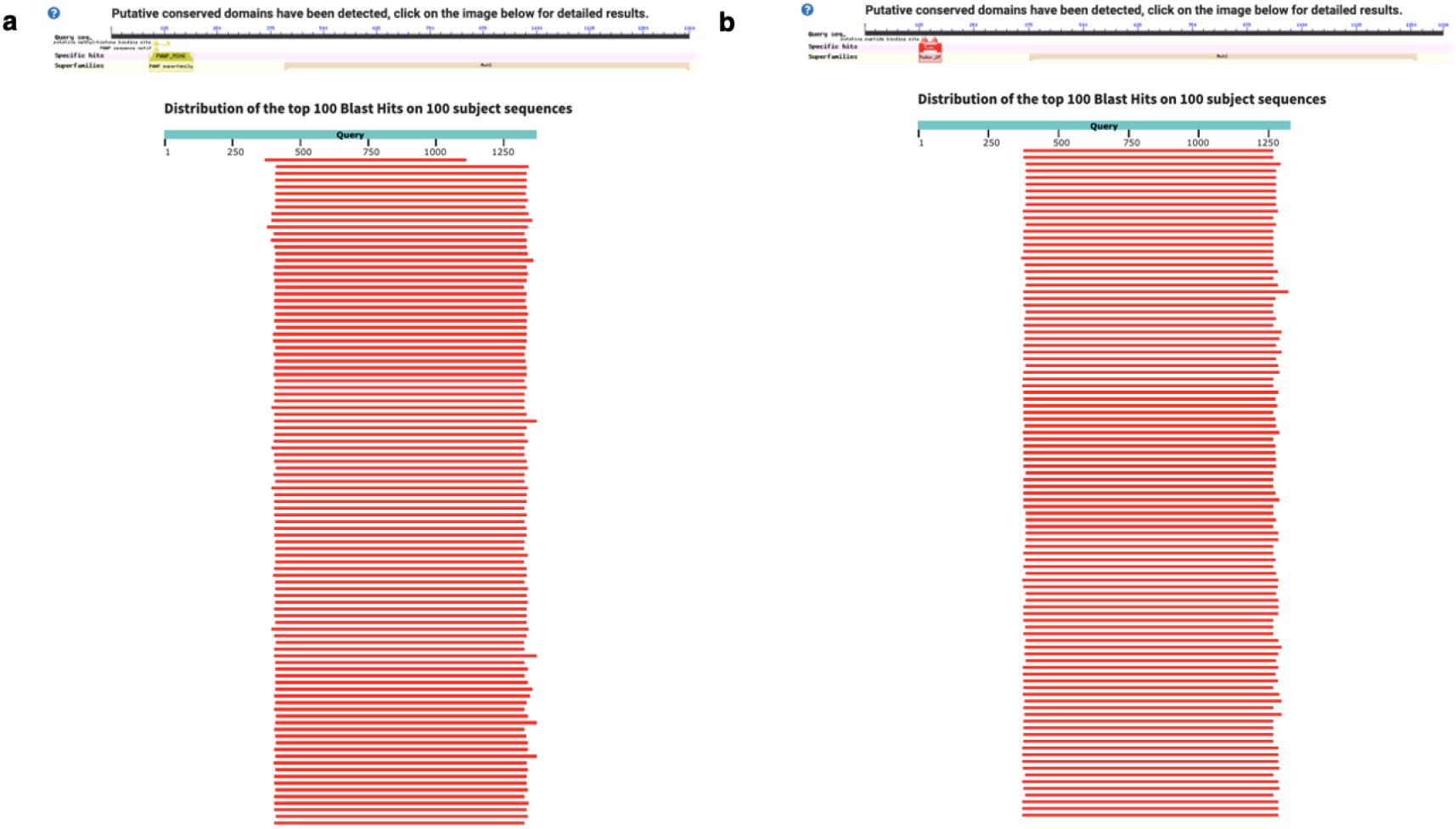
Results from MSH6 blastp queries in all archaeal NCBI proteomes. **a)** Human MSH6 with PWWP domain **b)** Arabidopsis MSH6 with Tudor domain. All results returned were annotated as MutS homologs. Top 100 shown from NCBI query results. Queries for Human MSH6 PWWP and Arabidopsis MSH6 Tudor did not yield any hits. In contrast, significant *blastp* results for PWWP and Tudor were found in MSH6 orthologs of distantly related eukaryotes, Cnidaria and Stramenopiles, respectively.

**Figure S4.**
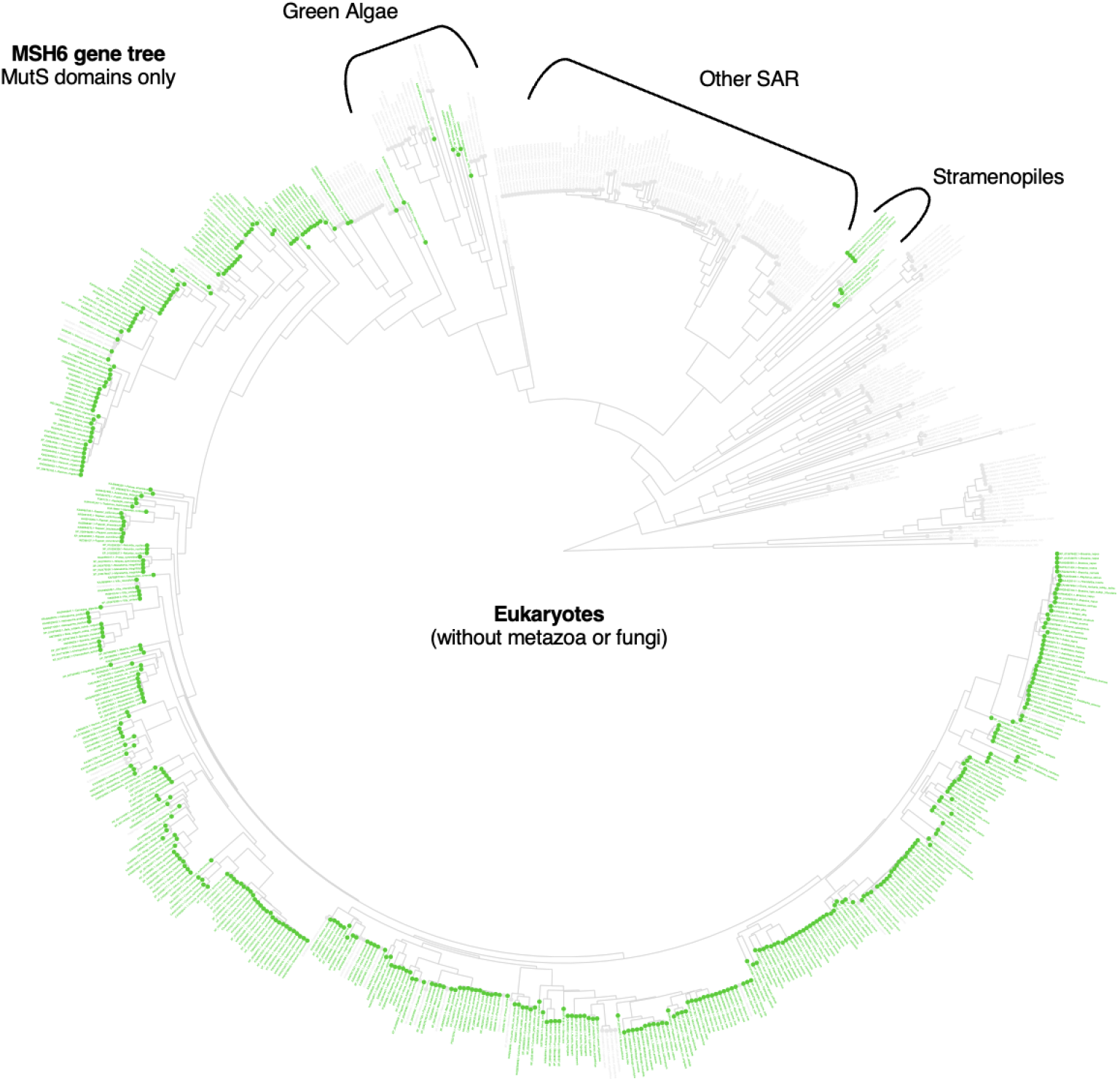
Lack of evidence that brown algae (Stramenopiles) acquired MSH6-Tudor via horizontal gene transfer from algae. Maximum likelihood tree from MutS domains of putative MSH6 orthologs in non-metazoan and non-fungal organisms. ML tree was completed with IQ-TREE from protein alignments of MSH6 only for domains MutS domain I, II, III, IV and V. Tips highlighted in green have *blastp* results for Tudor domain. The topology shows that the Stramenopiles’ MutS domains are sister to other SAR organisms, which lack Tudor domains, rather than sister to green algae which would have been a putative donor through horizontal gene transfer.

**Figure S5.**
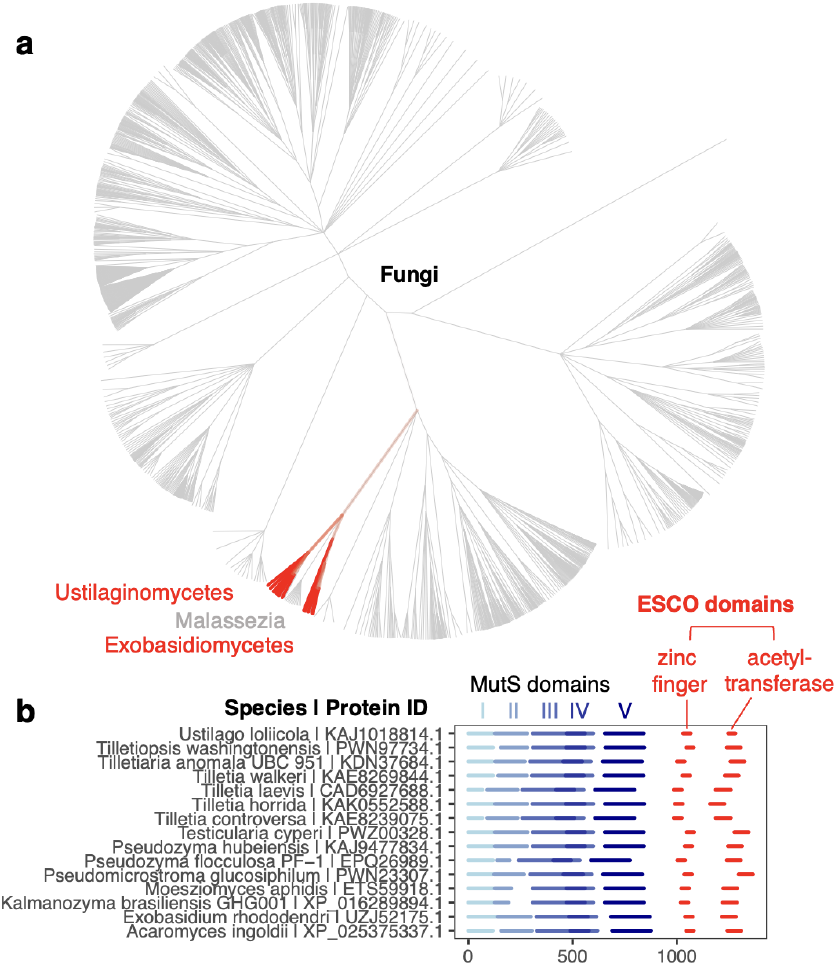
ESCO fusions to fungal MSH6. **a)** NCBI taxonomy tree of fungi with branches in red marking presence of ESCO domains, **b)** Domain architectures of MSH6-ESCO, aligned to MutS domain I.

**Figure S6.**
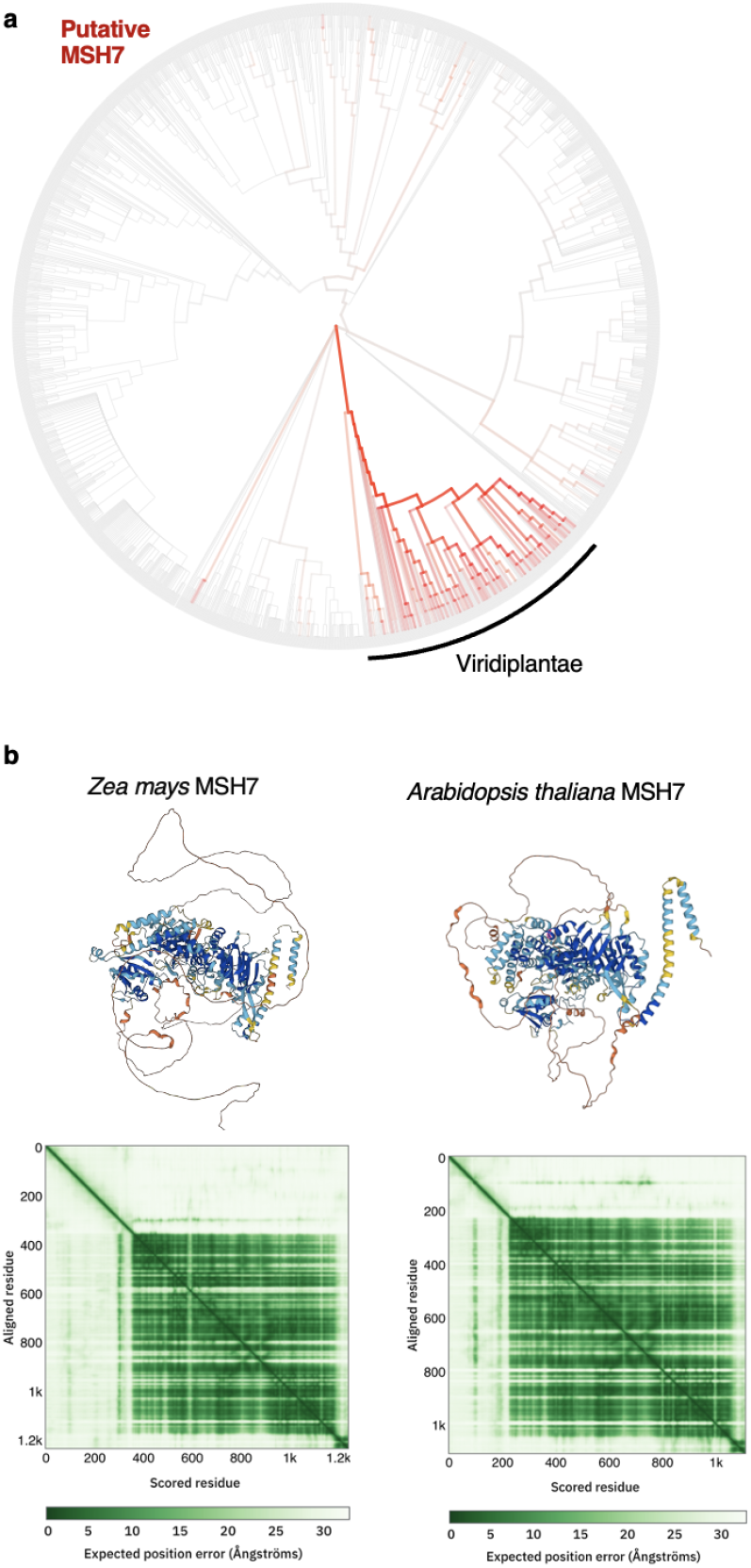
MSH7 in plants lacks a Tudor domain. **a)** Eukaryotes with NCBI branches colored (red) according to fraction with predicted MSH7 orthologs. 274/314 Viridiplantae species had predicted MSH7 orthologs, with 0 showing evidence of Tudor domains, **b)** Illustrative examples from *Zea mays* and *Arabidopsis thaliana*. Alphafold2 predicted structures of MutS homolog 7 (MSH7) in plants. MSH7 arose in plants via duplication of MSH6. The lack of histone domain at the terminus of MSH7 is also confirmed by domain prediction with Interproscan.

**Figure S7.**
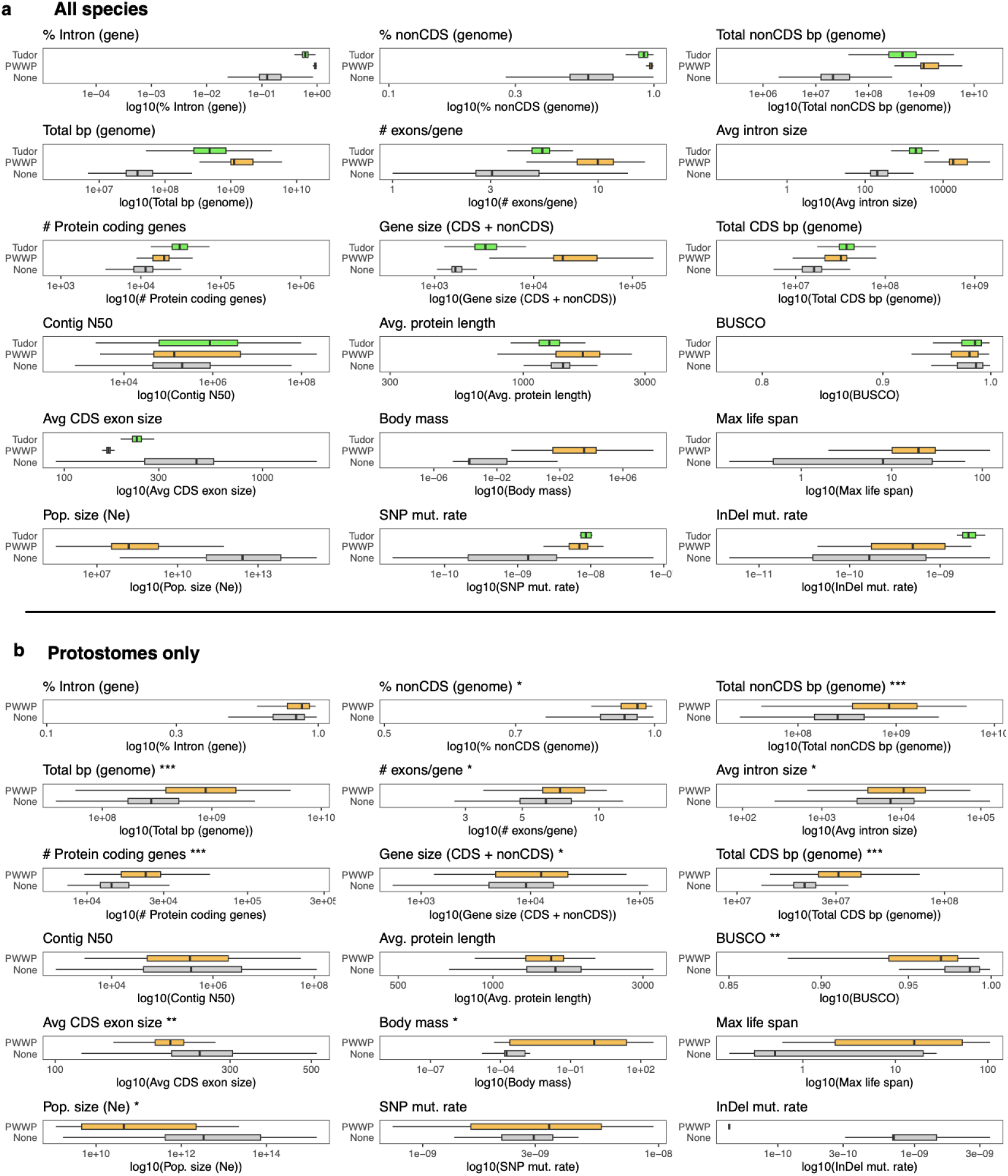
Species with histone readers fused to MSH6 differ in genomic and other traits. Summary of genomic and other characteristics among species with MSH6-Tudor, -PWWP, or no reader, Iog10 transformed values. Boxplots: box = median± interquartile range (IQR), whiskers = 1.5 x IQR. **a)** All species. Significance of differences reported in Fig. 4b,d. **b)** Only protestóme animals. Significance of difference from binomial regression: ***p<1×10^8^ **p<1×10’^3^, *p<0.05.

**Figure S8.**
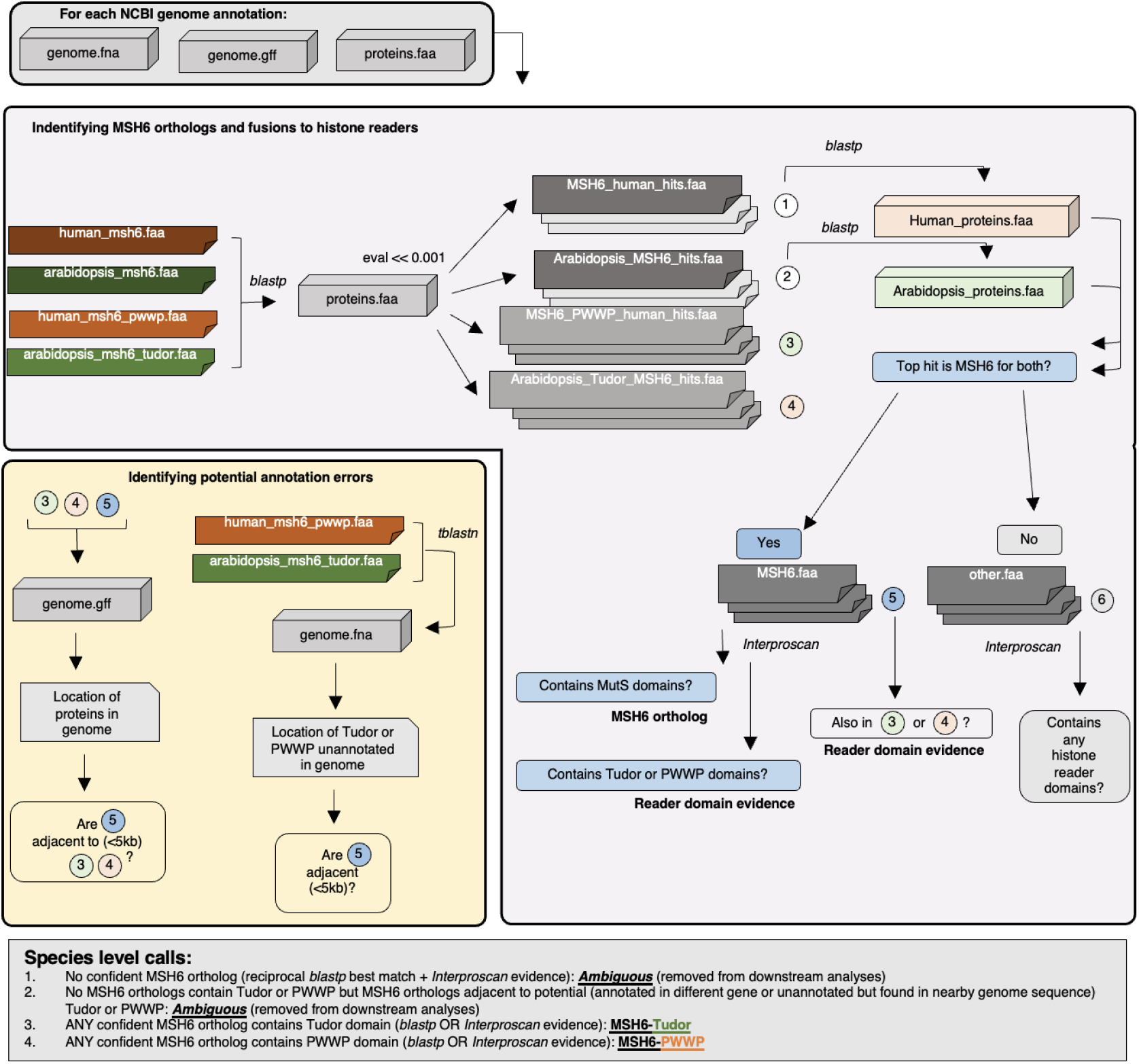
Workflow to identify MHS6 orthologs and their fusions to PWWP or Tudor domain histone readers.

## References

1. Manova, V. & Gruszka, D. DNA damage and repair in plants - from models to crops. Front. Plant Sci. 6, 885 (2015).

2. Nisa, M.-U., Huang, Y., Benhamed, M. & Raynaud, C. The Plant DNA Damage Response: Signaling Pathways Leading to Growth Inhibition and Putative Role in Response to Stress Conditions. Front. Plant Sci. 10, 653 (2019).

3. Yousefzadeh, M. et al. DNA damage—how and why we age? Elife 10, e62852 (2021).

4. Alberts, B. et al. Molecular Biology of the Cell. (Garland Science, 2002).

5. Friedberg, E. C. DNA Repair and Mutagenesis. Wiley & Sons, Limited, John, 2014).

6. Lindahl, T. Instability and decay of the primary structure of DNA. Nature 362, 709–715 (1993).

7. Sander, M. et al. Proceedings of a workshop on DNA adducts: Biological significance and applications to risk assessment Washington, DC, April 13–14, 2004. Toxicol. Appl. Pharmacol. 208, 1–20 (2005).

8. Sears, C. R. & Turchi, J. J. Complex Cisplatin-Double Strand Break (DSB) Lesions Directly Impair Cellular Non-Homologous End-Joining (NHEJ) Independent of Downstream Damage Response (DDR) Pathways *. J. Biol. Chem. 287, 24263–24272 (2012).

9. Mouret, S. et al. Cyclobutane pyrimidine dimers are predominant DNA lesions in whole human skin exposed to UVA radiation. Proc. Natl. Acad. Sci. U. S. A. 103, 13765–13770 (2006).

10. Lynch, M. Evolution of the mutation rate. Trends Genet. 26, 345–352 (2010).

11. Martincorena, I. & Luscombe, N. M. Non-random mutation: the evolution of targeted hypermutation and hypomutation. Bioessays 35, 123–130 (2013).

12. Lynch, M. et al. Genetic drift, selection and the evolution of the mutation rate. Nat. Rev. Genet. 17, 704–714 (2016).

13. Lynch, M. et al. The divergence of mutation rates and spectra across the Tree of Life. EMBO Rep. 24, e57561 (2023).

14. Li, F. et al. The Histone Mark H3K36me3 Regulates Human DNA Mismatch Repair through Its Interaction with MutSα. Cell 153, 590–600 (2013).

15. Huang, Y., Gu, L. & Li, G.-M. H3K36me3-mediated mismatch repair preferentially protects actively transcribed genes from mutation. J. Biol. Chem. 293, 7811–7823 (2018).

16. Kolasinska-Zwierz, P. et al. Differential chromatin marking of introns and expressed exons by H3K36me3. Nat. Genet. 41, 376–381 (2009).

17. Supek, F. & Lehner, B. Clustered Mutation Signatures Reveal that Error-Prone DNA Repair Targets Mutations to Active Genes. Cell 170, 534–547.e23 (2017).

18. Supek, F. & Lehner, B. Differential DNA mismatch repair underlies mutation rate variation across the human genome. Nature 521, 81–84 (2015).

19. Fang, H. et al. Deficiency of replication-independent DNA mismatch repair drives a 5-methylcytosine deamination mutational signature in cancer. Sci Adv 7, eabg4398 (2021).

20. Cavassim, M. I. A. et al. PRDM9 losses in vertebrates are coupled to those of paralogs ZCWPW1 and ZCWPW2. Proceedings of the National Academy of Sciences 119, e2114401119 (2022).

21. Huang, T. et al. The histone modification reader ZCWPW1 links histone methylation to PRDM9-induced double-strand break repair. Elife 9, e53459 (2020).

22. Wells, D. et al. ZCWPW1 is recruited to recombination hotspots by PRDM9 and is essential for meiotic double strand break repair. Elife 9, e53392 (2020).

23. Pradeepa, M. M., Sutherland, H. G., Ule, J., Grimes, G. R. & Bickmore, W. A. Psip1/Ledgf p52 binds methylated histone H3K36 and splicing factors and contributes to the regulation of alternative splicing. PLoS Genet. 8, e1002717 (2012).

24. Daugaard, M. et al. LEDGF (p75) promotes DNA-end resection and homologous recombination. Nat. Struct. Mol. Biol. 19, 803–810 (2012).

25. Sharma, S. et al. Affinity switching of the LEDGF/p75 IBD interactome is governed by kinase-dependent phosphorylation. Proceedings of the National Academy of Sciences 115, E7053–E7062 (2018).

26. Mahgoub, M. et al. Dual histone methyl reader ZCWPW1 facilitates repair of meiotic double strand breaks in male mice. Elife 9, e53360 (2020).

27. Eidahl, J. O. et al. Structural basis for high-affinity binding of LEDGF PWWP to mononucleosomes. Nucleic Acids Res. 41, 3924–3936 (2013).

28. Sundarraj, J., Taylor, G. C. A., von Kriegsheim, A. & Pradeepa, M. M. H3K36me3 and PSIP1/LEDGF associate with several DNA repair proteins, suggesting their role in efficient DNA repair at actively transcribing loci. Wellcome Open Res 2, 83 (2017).

29. Frigola, J. et al. Reduced mutation rate in exons due to differential mismatch repair. Nat. Genet. 49, 1684–1692 (2017).

30. Boukas, L., Razi, A., Bjornsson, H. T. & Hansen, K. D. Natural selection acts on epigenetic marks. bioRxiv 2020.07.04.187880 (2022) doi:10.1101/2020.07.04.187880.

31. Schuster-Böckler, B. & Lehner, B. Chromatin organization is a major influence on regional mutation rates in human cancer cells. Nature 488, 504–507 (2012).

32. Supek, F. & Lehner, B. Scales and mechanisms of somatic mutation rate variation across the human genome. DNA Repair 81, 102647 (2019).

33. Sun, Z. et al. H3K36me3, message from chromatin to DNA damage repair. Cell Biosci. 10, 9 (2020).

34. Chen, E. et al. Decorating chromatin for enhanced genome editing using CRISPR-Cas9. Proceedings of the National Academy of Sciences 119, e2204259119 (2022).

35. Hofstatter, P. G. & Lahr, D. J. G. Complex Evolution of the Mismatch Repair System in Eukaryotes is Illuminated by Novel Archaeal Genomes. J. Mol. Evol. 89, 12–18 (2021).

36. Laguri, C. et al. Human mismatch repair protein MSH6 contains a PWWP domain that targets double stranded DNA. Biochemistry 47, 6199–6207 (2008).

37. Li, F. et al. The histone mark H3K36me3 regulates human DNA mismatch repair through its interaction with MutSα. Cell 153, 590–600 (2013).

38. Huang, Y., Gu, L. & Li, G.-M. H3K36me3-mediated mismatch repair preferentially protects actively transcribed genes from mutation. J. Biol. Chem. 293, 7811–7823 (2018).

39. Moore, L. et al. The mutational landscape of human somatic and germline cells. Nature (2021) doi:10.1038/s41586-021-03822-7.

40. Rheinbay, E. et al. Analyses of non-coding somatic drivers in 2,658 cancer whole genomes. Nature 578, 102–111 (2020).

41. Zhang, X., Bernatavichute, Y. V., Cokus, S., Pellegrini, M. & Jacobsen, S. E. Genome-wide analysis of mono-, di- and trimethylation of histone H3 lysine 4 in Arabidopsis thaliana. Genome Biol. 10, R62 (2009).

42. Inagaki, S. et al. Gene-body chromatin modification dynamics mediate epigenome differentiation in Arabidopsis. EMBO J. 36, 970–980 (2017).

43. Liu, Y. et al. H3K4me2 functions as a repressive epigenetic mark in plants. Epigenetics Chromatin 12, 40 (2019).

44. Lu, Z. et al. The prevalence, evolution and chromatin signatures of plant regulatory elements. Nat Plants 5, 1250–1259 (2019).

45. Niu, Q. et al. A histone H3K4me1-specific binding protein is required for siRNA accumulation and DNA methylation at a subset of loci targeted by RNA-directed DNA methylation. Nat. Commun. 12, 3367 (2021).

46. Oya, S., Takahashi, M., Takashima, K., Kakutani, T. & Inagaki, S. Transcription-coupled and epigenome-encoded mechanisms direct H3K4 methylation. Nat. Commun. 13, 4521 (2022).

47. Mori, S. et al. Cotranscriptional demethylation induces global loss of H3K4me2 from active genes in Arabidopsis. bioRxiv 2023.02.17.528985 (2023) doi:10.1101/2023.02.17.528985.

48. Quiroz, D. et al. H3K4me1 recruits DNA repair proteins in plants. Plant Cell (2024) doi:10.1093/plcell/koae089.

49. Monroe, J. G. et al. Mutation bias reflects natural selection in Arabidopsis thaliana. Nature 602, 101–105 (2022).

50. Monroe, J. G. et al. Reply to: Re-evaluating evidence for adaptive mutation rate variation. Nature 619, E57–E60 (2023).

51. Kenchanmane Raju, S. K., Lensink, M., Kliebenstein, D. J., Niederhuth, C. & Monroe, G. Epigenomic divergence correlates with sequence polymorphism in Arabidopsis paralogs. New Phytol. (2023) doi:10.1111/nph.19227.

52. Ossowski, S. et al. The rate and molecular spectrum of spontaneous mutations in Arabidopsis thaliana. Science 327, 92–94 (2010).

53. Belfield, E. J. et al. Thermal stress accelerates Arabidopsis thaliana mutation rate. Genome Res. 31, 40–50 (2021).

54. Lu, Z. et al. Genome-wide DNA mutations in Arabidopsis plants after multigenerational exposure to high temperatures. Genome Biol. 22, 160 (2021).

55. Zhu, X. et al. Non-CG DNA methylation-deficiency mutations enhance mutagenesis rates during salt adaptation in cultured Arabidopsis cells. Stress Biology 1, 12 (2021).

56. Belfield, E. J. et al. DNA mismatch repair preferentially protects genes from mutation. Genome Res. 28, 66–74 (2018).

57. Yan, W., Deng, X. W., Yang, C. & Tang, X. The Genome-Wide EMS Mutagenesis Bias Correlates With Sequence Context and Chromatin Structure in Rice. Front. Plant Sci. 12, 579675 (2021).

58. Quiroz, D., Lensink, M., Kliebenstein, D. J. & Monroe, J. G. Causes of Mutation Rate Variability in Plant Genomes. Annu. Rev. Plant Biol. 74, 751–775 (2023).

59. Monroe, J. G. et al. Report of mutation biases mirroring selection in Arabidopsis thaliana unlikely to be entirely due to variant calling errors. bioRxiv 2022.08.21.504682 (2022) doi:10.1101/2022.08.21.504682.

60. Willing, E.-M. et al. UVR2 ensures transgenerational genome stability under simulated natural UV-B in Arabidopsis t haliana. Nat. Commun. 7, 1–9 (2016).

61. Quiroz, D. et al. The H3K4me1 histone mark recruits DNA repair to functionally constrained genomic regions in plants. bioRxiv 2022.05.28.493846 (2022).

62. Adé, J., Belzile, F., Philippe, H. & Doutriaux, M. P. Four mismatch repair paralogues coexist in Arabidopsis thaliana: AtMSH2, AtMSH3, AtMSH6-1 and AtMSH6-2. Mol. Gen. Genet. 262, 239–249 (1999).

63. Kolodner, R. Biochemistry and genetics of eukaryotic mismatch repair. Genes Dev. 10, 1433–1442 (1996).

64. Maurer-Stroh, S. et al. The Tudor domain ‘Royal Family’: Tudor, plant Agenet, Chromo, PWWP and MBT domains. Trends Biochem. Sci. 28, 69–74 (2003).

65. Grau-Bové, X. et al. A phylogenetic and proteomic reconstruction of eukaryotic chromatin evolution. Nat Ecol Evol 6, 1007–1023 (2022).

66. Lu, R. & Wang, G. G. Tudor: a versatile family of histone methylation ‘readers’. Trends Biochem. Sci. 38, 546–555 (2013).

67. Arendt, J. & Reznick, D. Convergence and parallelism reconsidered: what have we learned about the genetics of adaptation? Trends Ecol. Evol. 23, 26–32 (2008).

68. Wyder, S., Kriventseva, E. V., Schröder, R., Kadowaki, T. & Zdobnov, E. M. Quantification of ortholog losses in insects and vertebrates. Genome Biol. 8, R242 (2007).

69. Sekelsky, J. DNA Repair in Drosophila: Mutagens, Models, and Missing Genes. Genetics 205, 471–490 (2017).

70. Liu, H. & Zhang, J. Is the Mutation Rate Lower in Genomic Regions of Stronger Selective Constraints? Mol. Biol. Evol. 39, (2022).

71. Strenkert, D. et al. The landscape of Chlamydomonas histone H3 lysine 4 methylation reveals both constant features and dynamic changes during the diurnal cycle. Plant J. (2022) doi:10.1111/tpj.15948.

72. Curtis, B. A. et al. Algal genomes reveal evolutionary mosaicism and the fate of nucleomorphs. Nature 492, 59–65 (2012).

73. Wisecaver, J. H., Brosnahan, M. L. & Hackett, J. D. Horizontal gene transfer is a significant driver of gene innovation in dinoflagellates. Genome Biol. Evol. 5, 2368–2381 (2013).

74. Culligan, K. M. & Hays, J. B. Arabidopsis MutS homologs-AtMSH2, AtMSH3, AtMSH6, and a novel AtMSH7-form three distinct protein heterodimers with different specificities for mismatched DNA. Plant Cell 12, 991–1002 (2000).

75. Lin, Z., Nei, M. & Ma, H. The origins and early evolution of DNA mismatch repair genes--multiple horizontal gene transfers and co-evolution. Nucleic Acids Res. 35, 7591–7603 (2007).

76. Sloan, D. B. et al. Expansion of the MutS gene family in plants. bioRxivorg (2024) doi:10.1101/2024.07.17.603841.

77. Hou, F. & Zou, H. Two human orthologues of Eco1/Ctf7 acetyltransferases are both required for proper sister-chromatid cohesion. Mol. Biol. Cell 16, 3908–3918 (2005).

78. Campen, A. et al. TOP-IDP-scale: a new amino acid scale measuring propensity for intrinsic disorder. Protein Pept. Lett. 15, 956–963 (2008).

79. Lotthammer, J. M., Ginell, G. M., Griffith, D., Emenecker, R. J. & Holehouse, A. S. Direct prediction of intrinsically disordered protein conformational properties from sequence. Nat. Methods 21, 465–476 (2024).

80. Jack, B. R., Meyer, A. G., Echave, J. & Wilke, C. O. Functional Sites Induce Long-Range Evolutionary Constraints in Enzymes. PLoS Biol. 14, e1002452 (2016).

81. Rona, G. B., Eleutherio, E. C. A. & Pinheiro, A. S. PWWP domains and their modes of sensing DNA and histone methylated lysines. Biophys. Rev. 8, 63–74 (2016).

82. Sane, M., Diwan, G. D., Bhat, B. A., Wahl, L. M. & Agashe, D. Shifts in mutation spectra enhance access to beneficial mutations. Proc. Natl. Acad. Sci. U. S. A. 120, e2207355120 (2023).

83. Sane, M., Parveen, S. & Agashe, D. Mutation bias alters the distribution of fitness effects of mutations. bioRxiv 2024.03. 24.586369 (2024) doi:10.1101/2024.03.24.586369.

84. Sniegowski, P. D., Gerrish, P. J. & Lenski, R. E. Evolution of high mutation rates in experimental populations of E. coli. Nature 387, 703–705 (1997).

85. Carmel, L., Wolf, Y. I., Rogozin, I. B. & Koonin, E. V. Three distinct modes of intron dynamics in the evolution of eukaryotes. Genome Res. 17, 1034–1044 (2007).

86. Zhu, L. et al. Patterns of exon-intron architecture variation of genes in eukaryotic genomes. BMC Genomics 10, 47 (2009).

87. Buffalo, V. Quantifying the relationship between genetic diversity and population size suggests natural selection cannot explain Lewontin’s Paradox. Elife 10, e67509 (2021).

88. Wang, Y. & Obbard, D. J. Experimental estimates of germline mutation rate in eukaryotes: a phylogenetic meta-analysis. Evol Lett 7, 216–226 (2023).

89. Beichman, A. C., Zhu, L. & Harris, K. The Evolutionary Interplay of Somatic and Germline Mutation Rates. Annu Rev Biomed Data Sci (2024) doi:10.1146/annurev-biodatasci-102523-104225.

90. Jones, P. et al. InterProScan 5: genome-scale protein function classification. Bioinformatics 30, 1236–1240 (2014).

91. Liu, Y. et al. Expanded diversity of Asgard archaea and their relationships with eukaryotes. Nature 593, 553–557 (2021).

92. Yang, Z., Kumar, S. & Nei, M. A new method of inference of ancestral nucleotide and amino acid sequences. Genetics 141, 1641–1650 (1995).

93. Kumar, S. et al. TimeTree 5: An Expanded Resource for Species Divergence Times. Mol. Biol. Evol. 39, (2022).

94. Jumper, J. et al. Highly accurate protein structure prediction with AlphaFold. Nature 596, 583–589 (2021).

95. Rondelet, G., Dal Maso, T., Willems, L. & Wouters, J. Structural basis for recognition of histone H3K36me3 nucleosome by human de novo DNA methyltransferases 3A and 3B. J. Struct. Biol. 194, 357–367 (2016).

96. Marino, A., Debaecker, G., Fiston-Lavier, A.-S., Haudry, A. & Nabholz, B. Effective population size does not explain long-term variation in genome size and transposable element content in animals. eLife (2024) doi:10.7554/elife.100574.1.

97. Tacutu, R. et al. Human Ageing Genomic Resources: new and updated databases. Nucleic Acids Res. 46, D1083–D1090 (2018).

98. Ives, A. R. & Garland, T., Jr. Phylogenetic logistic regression for binary dependent variables. Syst. Biol. 59, 9–26 (2010).

99. Ives, A. R. & Garland, T. Phylogenetic Regression for Binary Dependent Variables. in Modern Phylogenetic Comparative Methods and Their Application in Evolutionary Biology: Concepts and Practice (ed. Garamszegi, L. Z.) 231–261 (Springer Berlin Heidelberg, Berlin, Heidelberg, 2014).

